# High-Throughput Screening Reveals Potential Inhibitors Targeting Trimethoprim-Resistant DfrA1 Protein in *Klebsiella pneumoniae* and *Escherichia coli*

**DOI:** 10.1101/2024.11.18.624070

**Authors:** Soharth Hasnat, Soaibur Rahman, Meherun Binta Alam, Farha Mohi Suin, Farzana Yeasmin, Tanjila Suha, Nahuna Tanjin Supty, Sal Sabila, Animesh Chowdhury, A.D.A. Shahinuzzaman, M Murshida Mahbub, M. Nazmul Hoque, Tofazzal Islam

## Abstract

The DfrA1 protein provides trimethoprim resistance in bacteria, especially *Klebsiella pneumoniae* and *Escherichia coli*, by modifying dihydrofolate reductase, which reduces the binding efficacy of the antibiotic. Thus, this study aimed to identify inhibitors of the trimethoprim-resistant DfrA1 protein through high-throughput computational screening of 3,601 newly synthesized chemical compounds sourced from the [ChemDiv](https://www.chemdiv.com/) database. We conducted high-throughput computational optimization and screening of a library containing 3,601 compounds against the DfrA1 protein from *K. pneumoniae* and *E. coli* to identify potential drug candidates (DCs). Through this extensive approach, we identified six promising DCs, labeled DC1 to DC6, as potential inhibitors of DfrA1. Each DC demonstrated strong initial binding affinity and favorable chemical interactions with the DfrA1 binding sites when compared to the effective drug Iclaprim (effective antibiotic against DfrA1), used as a control. To validate these findings, we further investigated the molecular mechanisms of inhibition, focusing on the thermodynamic properties of the promising DCs. Furthermore, molecular dynamics simulation (MDS) validated the inhibitory efficacy of these six DCs against the DfrA1 protein. Our results showed that DC4 (an organoflourinated compound) and DC6 (a benzimidazol compound) showed superior efficacy against the DfrA1 protein than the control drug, particularly regarding stability, solvent-accessible surface area, solvent exposure, polarity, and binding site interactions, which influence their residence time and efficacy. Overall, findings of this study suggest that DC4 and DC6 have the potential to act as inhibitors against the DfrA1, offering promising prospects for the treatment and management of infections caused by trimethoprim-resistant *K. pneumoniae* and *E. coli* in both humans and animals.

## Introduction

Inventions of new chemical compounds against antibiotic-resistant pathogens are the critical demand for world health security. Computational chemistry plays a central role in many drug discovery applications as computational methods help understand the in-sight of physicochemical properties of small-molecule drugs ^1^. Optimizing a vast array of organic compounds, particularly hit-to-lead candidates, has significantly enhanced drug-target binding affinity. Large-scale screening, powered by supercomputing, offers new hope in addressing the growing challenge of antibiotic resistance ^2^. The creation of a computational library containing diverse organic compounds, enriched with elements like oxygen, nitrogen, fluorine, sulfur, and chlorine, makes drug discovery more comprehensive and sophisticated ^3,4^. The presence of charged electrostatic regions with electronegative atoms facilitates nucleophile attraction, a critical factor in drug-protein interactions ^5^. *In-silico* techniques, such as drug-receptor interaction analysis and chemical attraction identification, now enable precise predictions before *in-vivo* experiments. Antimicrobial resistance (AMR) has emerged as a global public health crisis, threatening the efficacy of antibiotics against bacterial infections ^6^. The antibiotic trimethoprim (TMP) is commonly used to treat various bacterial infections; however, its effectiveness is often compromised by the rapid emergence of TMP-resistant bacterial strains ^7,8^. Recent studies reveal a widespread prevalence of resistance to TMP in infections caused by *Escherichia coli* and *Klebsiella* species, posing significant challenges for effective treatment ^7,9^. TMP mediates its antibacterial effect by targeting bacterial dihydrofolate reductase (DfrA1), a ubiquitous enzyme found in a large group of bacteria including the members of Gram-negative *Enterobacteriaceae* ^7,10^. Pathogenic bacteria such as *E. coli*, *K. pneumoniae*, *Pseudomonas aeruginosa* etc. are rapidly evolving resistance mechanisms against TMP, rendering standard treatments increasingly ineffective ^7,9,11^. The surge in TMP resistance leads to prolonged illnesses, higher healthcare costs, increased mortality, and a greater risk of spreading infections, underscoring the urgent need for innovative solutions ^12^. According to the Centers for Disease Control and Prevention, TMP-resistant bacteria account for more than 9,000 healthcare-associated infections each year in the United States ^13^. A significant contributor to this resistance is the widespread occurrence of the *DfrA1* gene, which can confer a 190 to 1,000-fold increase in TMP resistance ^14^. In light of the challenges posed by TMP-resistant bacteria, particularly *E. coli* and *K. pneumoniae*, our research aims to identify novel promising drug candidates (DCs) by targeting the highly virulent DfrA1 protein, a major factor driving TMP resistance in these pathogens.

Computational tools like Gaussian offer a cost-effective and efficient approach to drug design and development, providing significant insights within a limited timeframe ^15,16^. These methods enable the prediction of drug geometries, structures, and physicochemical and biological properties. Density functional theory (DFT) technique, particularly when combined with cluster-based supercomputing, allows for the extraction of highly accurate information on biological systems in a short period ^17^. The effectiveness of drug-target interactions can also be anticipated by evaluating binding sites. Furthermore, DFT studies on the Highest Occupied Molecular Orbital (HOMO) and Lowest Unoccupied Molecular Orbital (LUMO) and electronic spectra provide valuable insights into molecular stability and reactivity ^18^. The use of computational platforms like Maestro (Schrödinger LLC) further enhances the drug discovery process, making it more cost-effective by revealing the stability of potential drug candidates in their interactions with receptor proteins ^19^. However, despite its value, this approach can sometimes be limited in accurately capturing the inherent reactivity of a molecule within a protein complex, such as the drug-binding pocket of a protein ^20^. Chemical compounds interact with the amino acids in these binding pockets, primarily through electrophilic or nucleophilic attraction to the side chain atoms of the amino acids ^19^. The computer-based software used in these analyses is often developed from lab-based experiments and incorporates various algorithmic calculations to minimize errors ^21^. Despite these advancements, additional methods are needed in drug discovery and development. For instance, Molecular Mechanics/Generalized Born Surface Area (MM/GBSA) calculations are frequently used to estimate the free energy of binding between a ligand and a receptor or protein during molecular dynamics simulations ^22^. This approach provides a more accurate computational prediction of receptor-ligand interactions.

In this study, we screened 3,601 newly synthesized chemical compounds containing various electronegative chemical species against the DfrA1 protein. While TMP is entirely ineffective against DfrA1, Iclaprim has been identified as the most effective drug targeting this virulent protein [23]. However, resistance has recently emerged, leading to a decrease in the effectiveness of Iclaprim by 60- to 700-fold ^23^. Given the emerging issue of AMR against DfrA1, particularly in the absence of a fully effective drug, there is an urgent need to identify new therapeutic agents to combat resistant pathogens. Thus, the identification of two drug candidates, DC4 and DC6, through the large-scale screening of a library of 3,601 chemical compounds, presents a promising opportunity for developing therapeutics targeting trimethoprim-resistant DfrA1 protein in *K. pneumoniae* and *E. coli*. This discovery opens new avenues for further wet lab experiments for validation of our computational findings.

## Results

### Protein-DCs binding simulations identify effective inhibitors of DfrA1

A comprehensive screening of 3,601 optimized compounds (**File S1**) against TMP-resistant DfrA1 protein identified six compounds (DC1 – DC6) as effective binders. These six DCs were highlighted as the promising inhibitors of DfrA1 due to their strong interaction profiles, which included H-bonds and other non-covalent interactions, as well as their high affinity for the drug-binding pockets. Remarkably, the binding free energies of these inhibitors were all lower than −9 kcal/mol (**Table 1**), indicating a strong binding potential. The analysis of the chemical structures of the DCs revealed the presence of charged atoms, such as oxygen and nitrogen, in the DC inhibitors. The chemical structure, IUPAC name, and SMILES notation of the most promising compounds, as well as the control, are presented in **Table 1**. The identification of the drug-binding pocket regions in DfrA1 protein showed that the binding pocket is centrally located within the protein, as indicated by the red coloration (**Fig. S1**). Further analysis of the screened compounds demonstrated that all six potential inhibitors (DCs), along with the control drug, bind within the identified pocket region (**Fig. 1**). During the initial screening, DC1 formed three H-bonds with the amino acids PRO18, ILE20, and SER49 within the binding pocket, with bond lengths ranging from 2.4 to 3.41 Å. Additionally, three non-covalent interactions were detected, with distances ranging from 2.27 to 5.15 Å (**Fig. 1A**). The electronegative atoms of DC2, primarily oxygen and nitrogen, formed four H-bonds with the amino acids PRO18, ILE20, GLN28, and SER49, with bond lengths ranging from 2.29 to 2.98 Å. Additionally, seven weak non-covalent interactions were identified between DC2 and the binding pocket amino acids of the DfrA1 (**Fig. 1B**). DC3 anchored to the binding pockets via three H-bonds and ten non-covalent interactions, including Pi-stacking, Pi-alkyl, Pi-sigma, and Van der Waals interactions, marking the highest number of non-covalent interactions among the six compounds and the control (**Fig. 1C**). The electronegative oxygen and nitrogen atoms in DC4 formed H-bonds with ILE14, ILE20, and SER49, with bond distances ranging from 2.14 to 3.20 Å, along with six non-covalent interactions within the complex (**Fig. 1D**). The pyridine-rich DC5 exhibited strong binding to the pocket amino acids primarily through non-covalent interactions, with GLU27, GLN28, and SER49 forming hydrogen bonds with DC5 at distances of 2.25 to 2.97 Å (**Fig. 1E**). DC6, characterized by its nitrogen-rich structure, demonstrated the second-highest number of non-covalent interactions within the pocket regions and formed three H-bonds with TRP22, GLU27, and GLN28, with bond lengths ranging from 2.48 to 2.83 Å (**Fig. 1F**). The control drug, Iclaprim, anchored within the binding pockets through two hydrogen bonds at distances of 2.3 to 2.77 Å and exhibited six non-covalent interactions. Comparative analysis showed that the six DCs demonstrated significantly higher affinities to the binding pockets than the control drug (**Fig. 1G**).

**Fig. 1.**
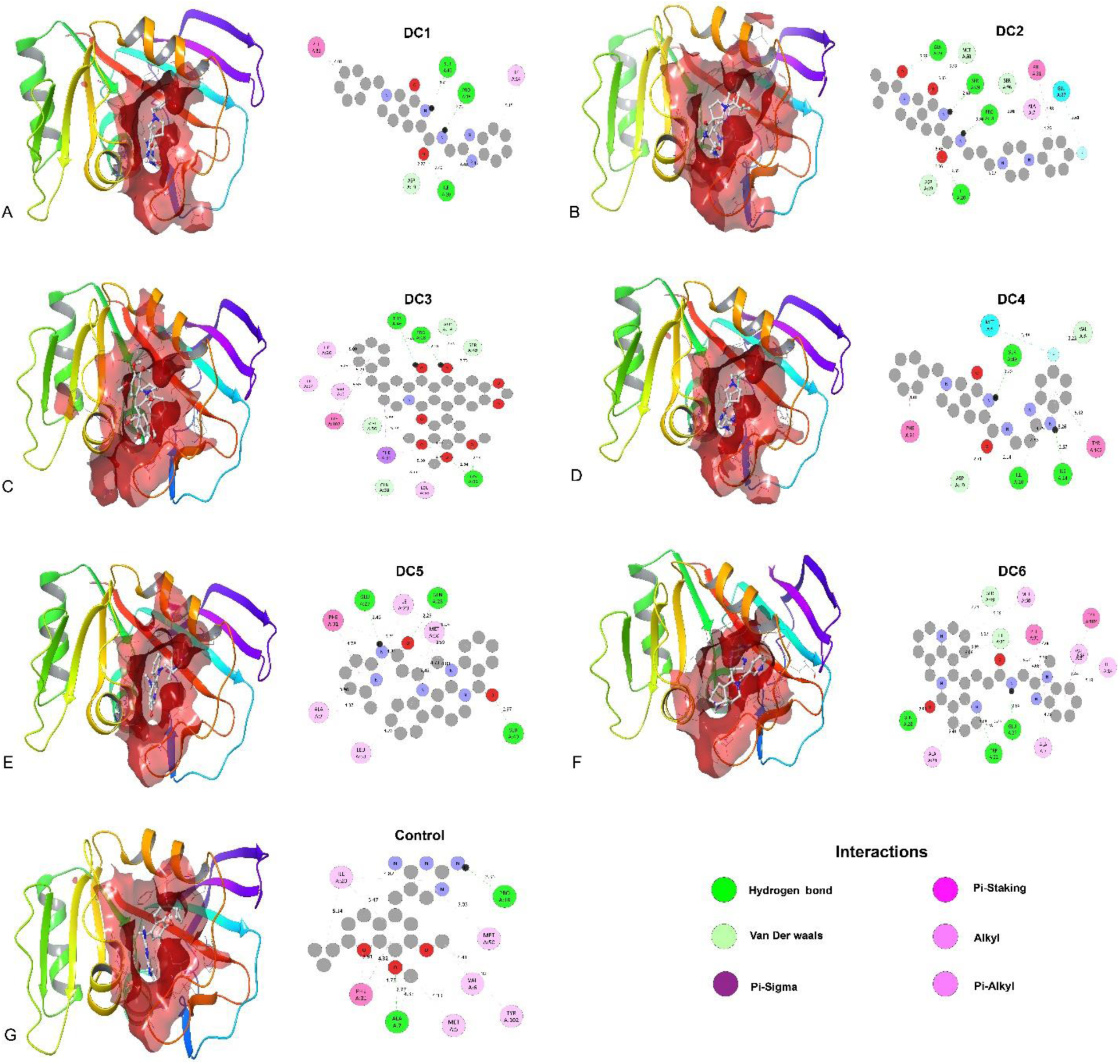
Interaction of identified potential drug candidates (DCs) with the binding pockets of the DfrA1 protein. The interaction diagrams with highlighted (red) pocket regions and binding details. Interaction of DCs with DfrA1 via pocket residues, showing binding distance A) DC1-DfrA1 complex. B) DC2-DfrA1 complex. C) DC3-DfrA1 complex. D) DC4-DfrA1 complex. E) DC5-DfrA1 complex. F) DC6-DfrA1 complex. G) Control-DfrA1 complex. The initial interaction and binding distance of each contact are presented in detail in each panel.

**Table 1:**
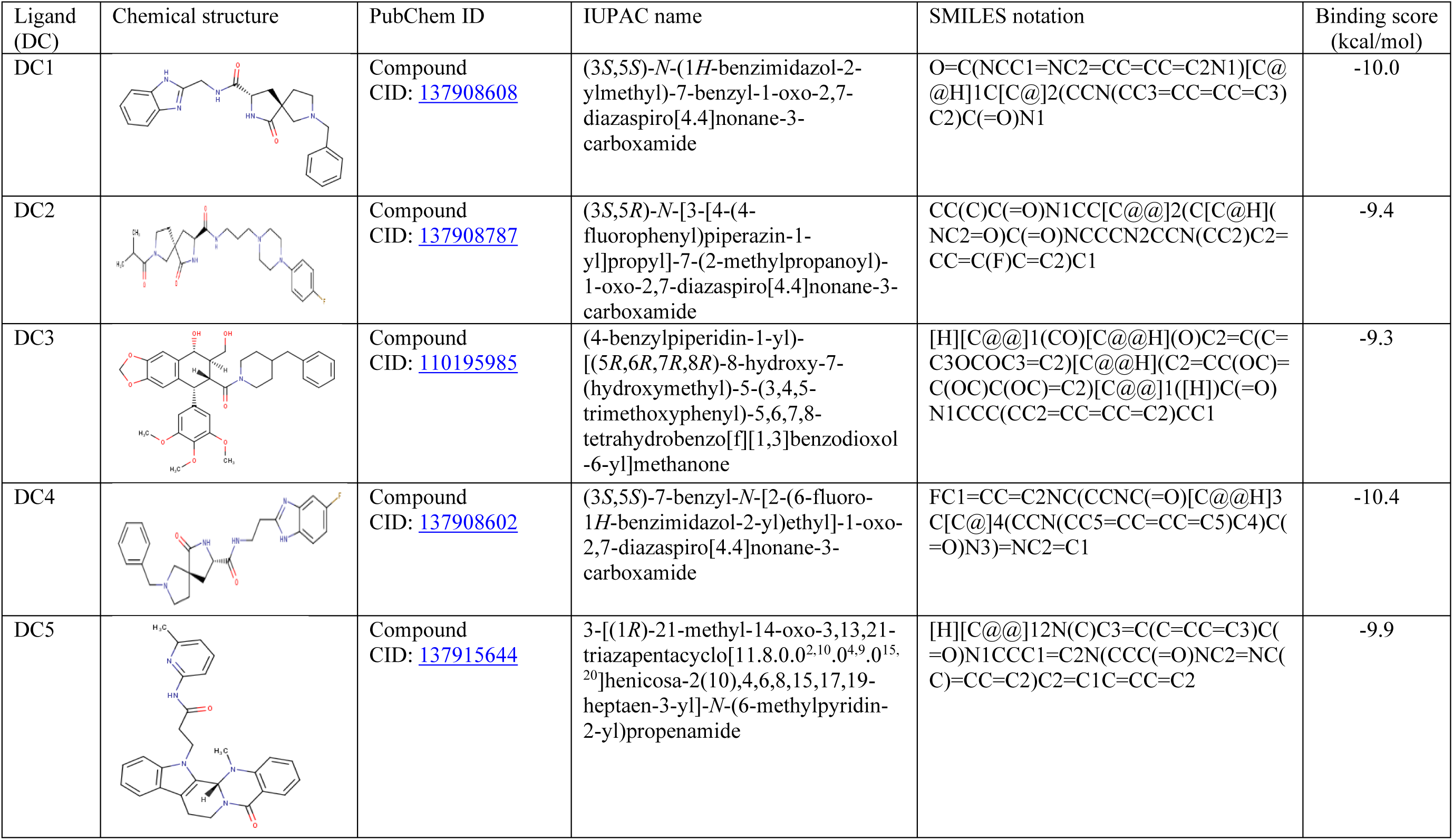
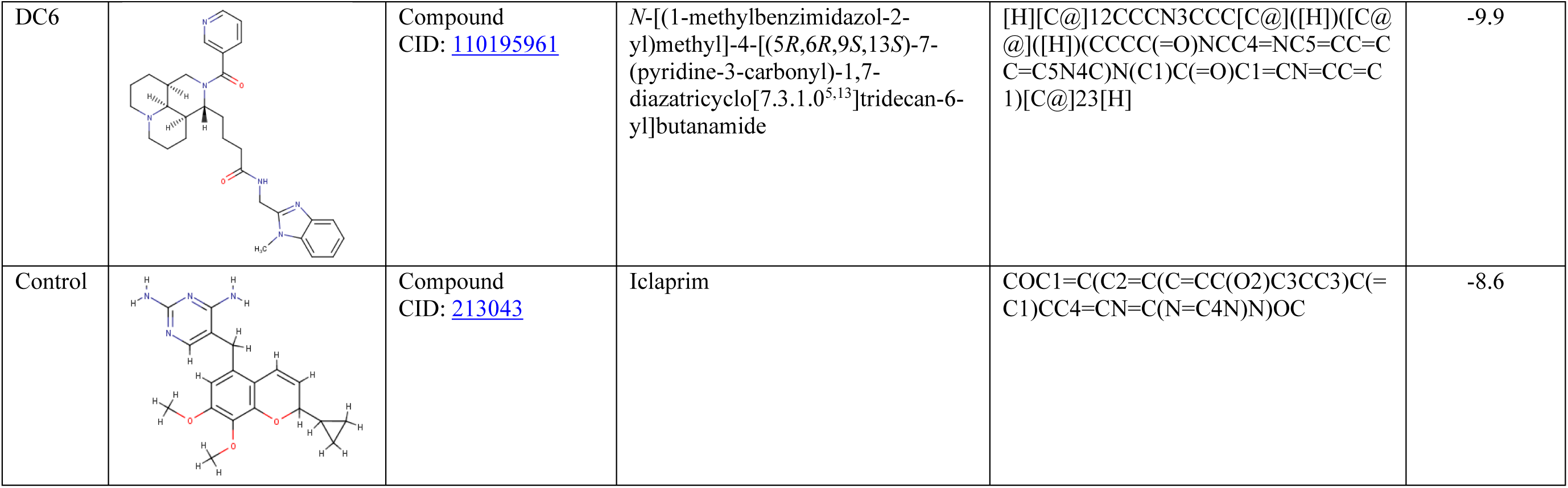
Chemical structures, annotation and docking scores of the best interacting drug candidates (DCs) and the control drug.

### Molecular orbital analysis highlights the potential of DCs as drug candidates against DfrA1

A key objective in investigating the molecular orbitals of potential DfrA1 protein inhibitors was to elucidate their electronic properties. Our analysis focused on the energy gap (ΔEHOMO-LUMO), which provides insights into the electronic distribution and reactivity of these compounds. For DC1, the HOMO and LUMO energy levels were determined to be −5.72 eV and −0.50 eV, respectively, yielding an energy gap (ΔEHOMO-LUMO) of 5.22 eV. The density of states (DOS) plot for DC2 revealed an energy gap (ΔEHOMO-LUMO) of 5.03 eV, which is lower than that of DC1. For DC3, the HOMO and LUMO energy levels were −5.68 eV and −0.22 eV, respectively, resulting in an energy gap (ΔEHOMO-LUMO) of 5.46 eV. According to the DOS plot for DC4, the energy gap (ΔEHOMO-LUMO) was 4.75 eV, with HOMO and LUMO energy levels at −5.59 eV and −0.84 eV, respectively. For DC5, the HOMO and LUMO energy levels were −5.46 eV and −1.05 eV, respectively, resulting in an energy gap (ΔEHOMO-LUMO) of 4.41 eV. DC6 exhibited the lowest energy gap (ΔEHOMO-LUMO) of 3.18 eV, indicating the smallest energy gap among the six compounds. In comparison, the DOS plot for the control compound showed a ΔEHOMO-LUMO of 4.77 eV. These ΔEHOMO-LUMO values suggest that some potential inhibitors are more reactive than the control drug, while others are less reactive (**Fig. 2**).

**Fig. 2.**
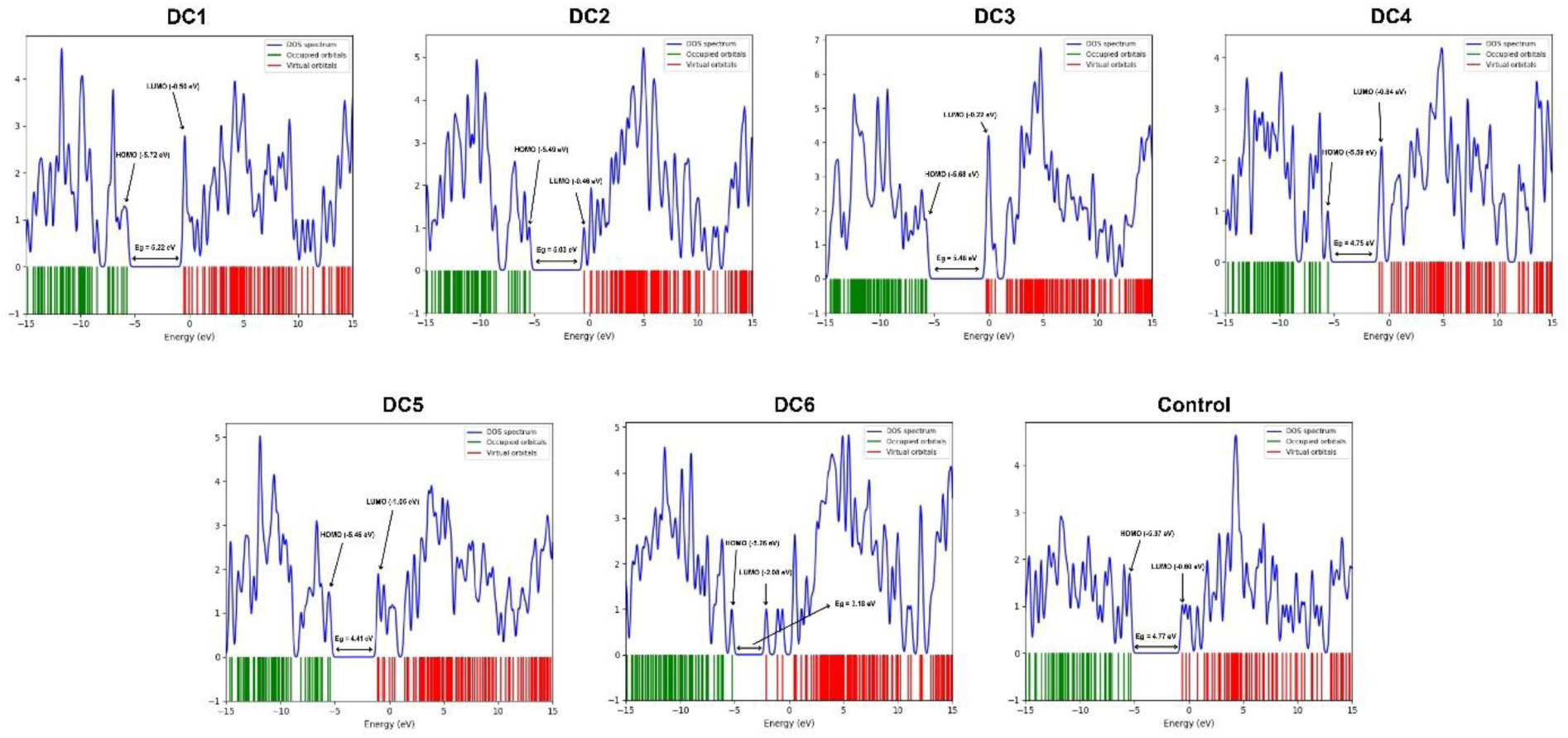
Density of state (DOS) plot of HOMO–LUMO and energy gap of best-interacting DCs along with the control. The green and red dots in the diagrams show HOMO and LUMO orbitals.

### Electronic absorption spectra (UV-Vis) provide insights into molecular electronic transitions of DCs

We then focused on the electronic transitions of the molecules using DFT methods. Calculations of electronic absorption spectra confirmed the maximum absorbances and excitation energies for six selected molecules, including the control (**Fig. S2**). Among the six molecules, DC6 exhibited the highest peak intensity at 401.90 nm in the S0 → S2 excited state, with an excitation energy of 3.0849 eV and configurations of (−0.11274) H-6→L, (0.25326) H-3→L, and (0.62777) H-2→L. DC4 and DC5 showed absorption bands for S0 → S1 at 303.34 nm and 332.43 nm, with energies of 4.0873 eV (configurations: (0.27669) H→L, (0.63590) H→L+1, (−0.11557) H→L+2) and 3.7297 eV (configurations: (−0.34646) H-1→L, (0.18515) H-1→L+1, (0.57349) H→L), respectively. DC1, DC2, and DC3 showed peaks at 278.86 nm, 257.30 nm, and 261.71 nm with excitation energies of 4.4460 eV, 4.8186 eV, and 4.7374 eV, respectively. The control drug, Iclaprim, displayed an absorption peak at 290.07 nm in the S0 → S3 transition with an excitation energy of 4.2743 eV (**Table S1**).

### ESP calculations enhance understanding of the DCs chemical reactivity

As the six compounds preferentially bind to the binding site amino acids of the DfrA1 protein, we proceeded to analyze the surface characteristics of each DC and the control drug. The ESP charge ranges for DC1 through DC6 were −0.142 to 0.142 a.u., −0.103 to 0.103 a.u., −0.109 to 0.109 a.u., −0.101 to 0.101 a.u., −0.127 to 0.127 a.u., and −0.141 to 0.141 a.u., respectively. In contrast, the control drug, Iclaprim, exhibited a charge range of −0.0592 to 0.0592 a.u., which was lower than that of all the studied DCs (**Fig. 3**). The surface analysis of the potential DCs revealed that highly electronegative atoms appeared as red zones, while electropositive regions were depicted in blue. Neutral atoms were represented by green zones, and slightly electronegative and positive atoms formed yellow and turquoise zones, respectively. Notably, the DC compounds did not display extreme positive or negative ESP values, suggesting that they are less likely to experience non-specific interactions or exhibit weak binding with the target. The ESP values for the six compounds were similar and indicated better performance compared to the control drug (**Fig. 3**). Comparing molecular interactions and ESP surfaces revealed that the charged surfaces of the compounds contributed to covalent interactions with the binding pocket amino acids, while partially charged surfaces were often involved in non-covalent interactions (**Figs. 1** and **3**).

**Fig. 3.**
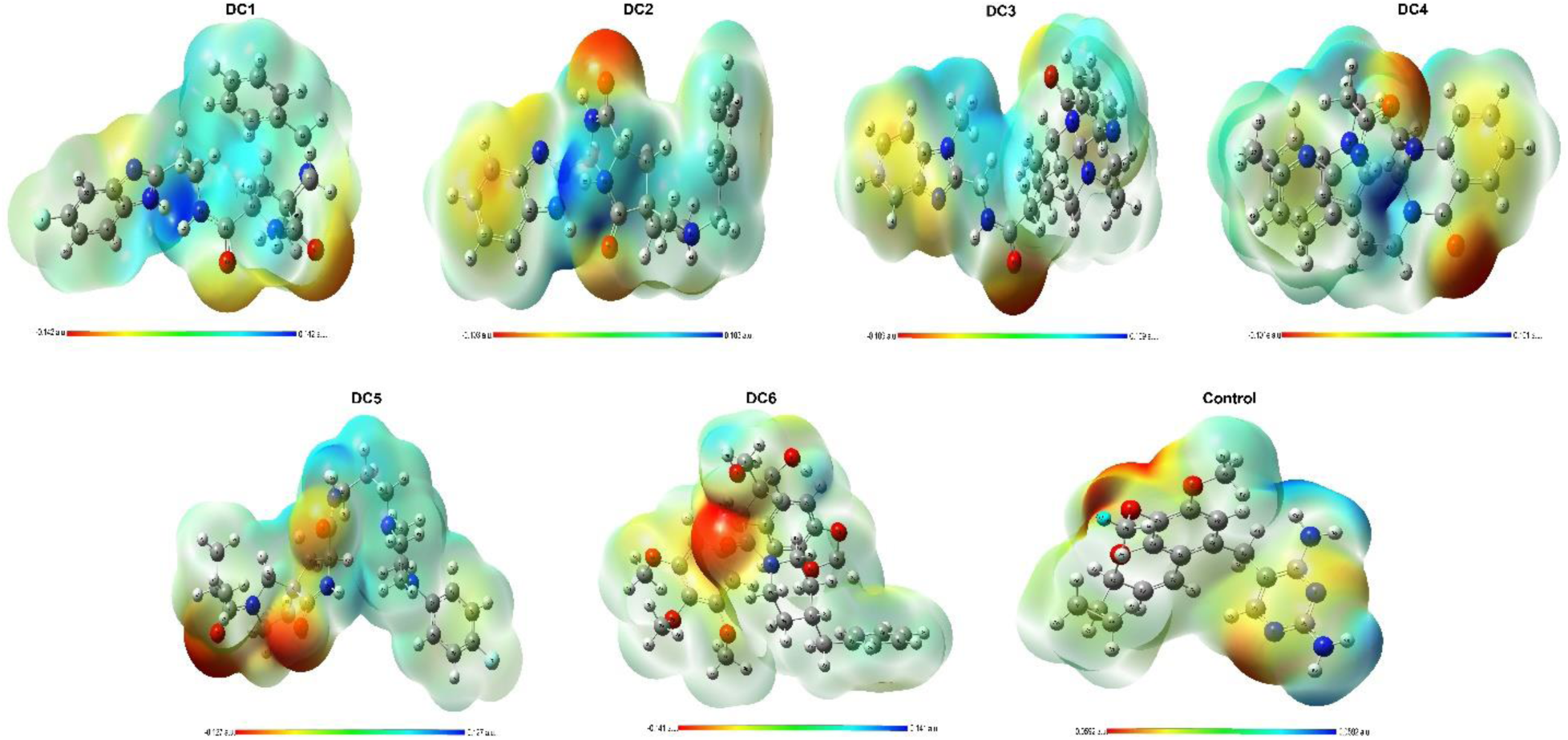
The electrostatic surface potential (ESP) map of six potential inhibitors of DfrA1 and a control. Different colors are used to specify the charge distribution, such as blue, red, and green colors, which are used to determine the positive charge, negative charge, and neutral charge, respectively. Short electron density and poor interaction are identified in the blue region, but the red region contains high electron density and potential interaction.

### MDS of the drug candidates detects stable interactions with DfrA1 protein

One critical aspect of identifying a novel drug compound targeting a specific protein involved observing stable interactions between the receptor and ligand throughout simulation trials. For DC1, two stable interactions were observed: one between two oxygen atoms of DC1 and amino acids THR46 and TYR58 through the H-bonding, and another H-bond was formed between SER96 and the hydrogens positively charged carbon of DC1 (**Fig. 4A**). However, MDS for the interaction of the DC2 with binding pocket regions of DfrA1 displayed no stable covalent interactions, besides interaction fractions diagram showed weaker non-covalent interactions (**Fig. 4B**). For DC3, two negatively charged oxygen atoms of DC1 formed stable bonds throughout the simulation trajectories with amino acids THR46 and SER96. The electro-negative oxygen of the ALA24 anchored to the positively charged hydrogen of DC3, which led to the formation of another H-bond (**Fig. 4C**). The MDS for the DC4 interaction with binding pocket regions of DfrA1 demonstrated the most robust attachment by forming four H-bonds (2 by electrophilic attack and two by nucleophilic attack) via PRO18, ILE20, THR46 and SER49. The reported interaction fraction was also relatively high compared to other atoms (**Fig. 4D**). The DC5 showed three H-bonds and one non-covalent interaction, where H-bonds formed via one nucleophilic and two electrophilic reactions, and other non-covalent interaction formed by the pyridine ring (**Fig. 4E**). Another strong interaction was observed between DC6 and four amino acids: ALA7, TRP22, ALA24, and GLY26. DC6 engaged with ALA7, TRP22, and GLY26 via its electronegative oxygen and nitrogen atoms, while the electronegative oxygen of ALA24 attracted the hydrogen atoms of DC6. These interactions further demonstrate the affinity of DC6 for the binding site, contributing to its potential effectiveness as a drug candidate (**Fig. 4F**). For the control compound, three covalent interactions were identified, showing strong binding within the DfrA1 binding pocket. ALA7 was attracted to the control compound through highly negative nitrogen atoms and an electronegative side chain, forming a H-bond. Additionally, the electronegative side chain of SER96 attracted the positively charged hydrogen atoms, leading to further bonding (**Fig. 4G**). These findings indicated that DC1, DC4, and DC6 exhibit robust interactions with the target binding regions, in some cases demonstrating even stronger interactions than the control compound, Iclaprim.

**Fig. 4.**
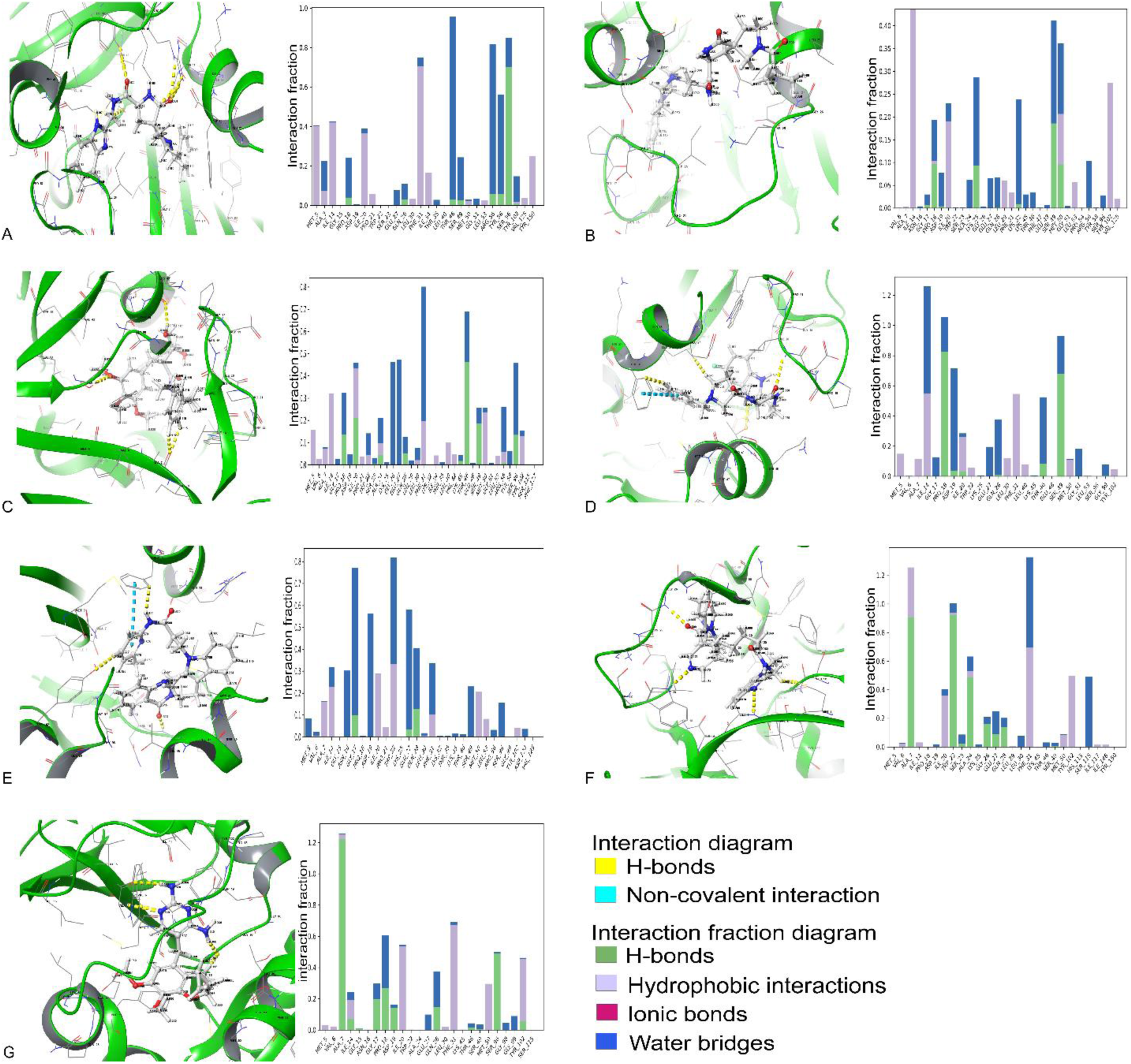
Stable binding contact of identified potential drug candidates (DCs) with the pocket’s residues of the DfrA1 protein. H-bonds and non-covalent interaction of DCs with DfrA1 via pocket residues and diagram of interaction strength A) DC1-DfrA1 complex. B) DC2-DfrA1 complex. C) DC3-DfrA1 complex. D) DC4-DfrA1 complex. E) DC5-DfrA1 complex. F) DC6-DfrA1 complex. G) Control-DfrA1 complex. The interaction contact and strength of each contact are presented in detail in each panel.

### MSD curves reveal the most potential inhibitors of the DfrA1 protein

Given the stable interactions observed with the DCs, we proceeded to analyze the Root Mean Square Deviation (RMSD) of the protein-ligand complexes during binding, as well as the Root Mean Square Fluctuation (RMSF) of the protein. The RMSD of the DfrA1 protein showed fluctuations of up to 2Å when interacting with all DCs, except DC1 and DC3. In contrast, the control compound exhibited fluctuations around 3.5Å, which is within an acceptable range. However, the lower RMSD values for the protein when bound to the DCs suggest that these compounds offer greater stability, making them more favorable than the control in terms of protein stability during interaction (**Fig. 5A**). The most significant aspect of the MDS was the ligand RMSD, crucial for assessing the strength and stability of the ligand within the binding pocket of DfrA1. The MDS results revealed that DC4 had the lowest RMSD value, consistently below 1Å, indicating strong stability during interaction. In contrast, DC2 exhibited the highest RMSD displacement, reaching nearly 6Å, suggesting instability when interacting with the binding pocket. Other DCs, as well as the control, exhibited displacements of up to 3Å, remaining within the acceptable stability threshold (**Fig. 5B**). The RMSF plots proved valuable for assessing local variations along the protein chain. Based on these plots, the amino acid regions at positions 20-35, 65-75, 120-125, and 145-150 displayed the highest fluctuations during the simulation compared to other regions. However, the reported fluctuations for these regions were approximately 3Å. The only anomaly was observed in the control compounds, which exhibited the highest displacement, around 5Å, in the 20-35 amino acid region. Additionally, higher RMSF values were noted for all other regions compared to the DCs (**Fig. 5C**).

**Fig. 5.**
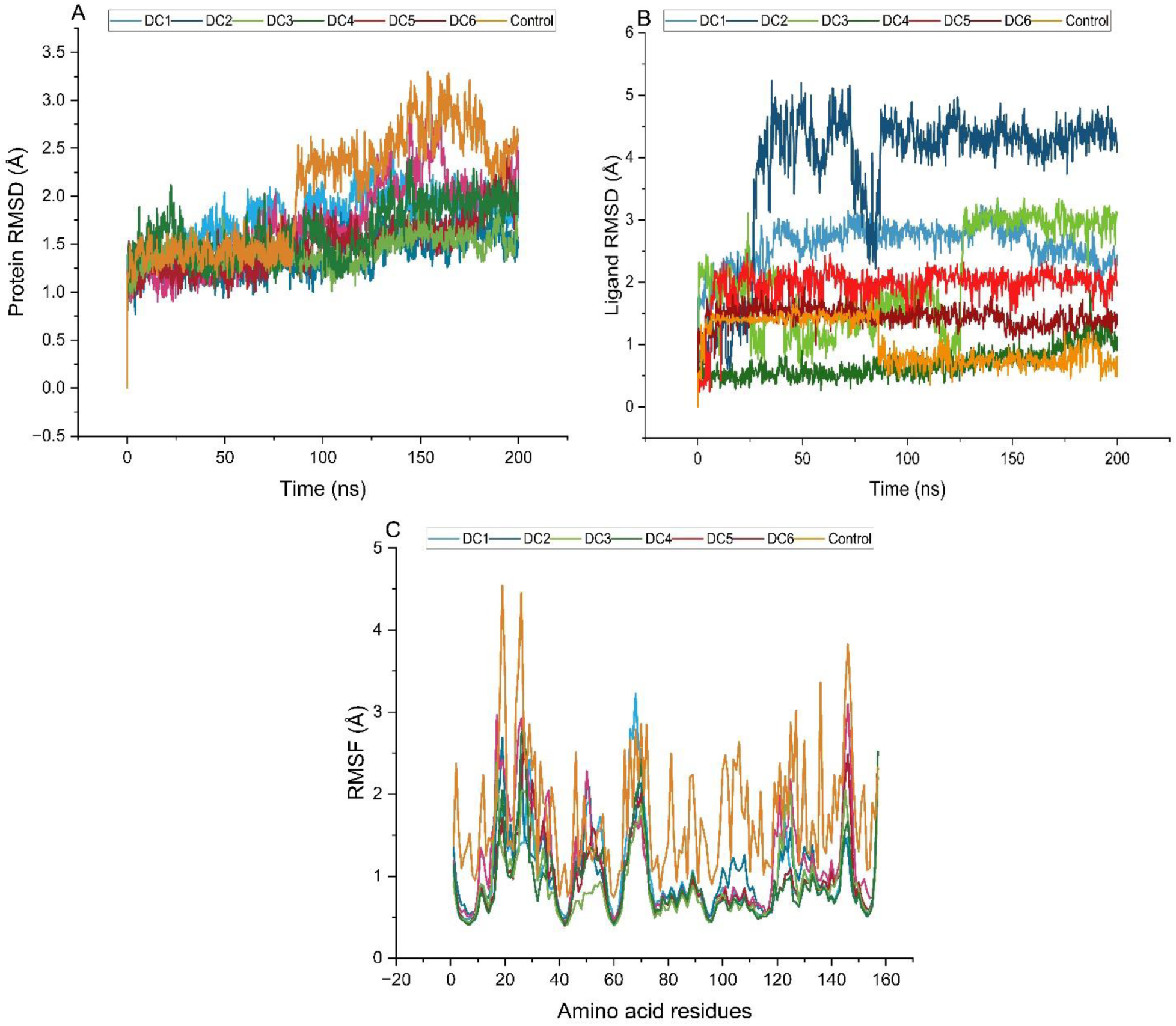
Thermodynamic stability analysis of the protein, potential drug candidates (DCs), and amino acids over 200 ns simulation trajectories. A) Overall fluctuations of DfrA1 during its interaction with DCs throughout the MDS period. B) Overall fluctuations of all DCs and a control drug during their interaction with DfrA1 in the MDS period. C) The RMSF curve shows the displacement of each amino acid throughout the MDS process.

### DCs properties analysis discloses the effectiveness as potential drug candidates

The drug-likeness properties, such as Molecular Surface Area (MolSA), were typically calculated based on the van der Waals surface of the molecule. The maximum MolSA value was recorded at 500Å^2^ for DC3, while the control compound showed the lowest value of around 325Å^2^. The MolSA values for the other five DCs ranged between 350 and 450Å^2^ (**Fig. 6A**). We then calculated the Solvent Accessible Surface Area (SASA) values for the DCs and the control drug. In MDS, SASA refers to the surface area of a compound that is accessible to solvent molecules, typically water. The results indicated that DC2 had the most accessible surface area, with SASA values ranging from 200Å^2^ to 400Å^2^. The second-highest SASA value was observed for DC3, ranging from 100Å^2^ to 250Å^2^. The SASA values for the other four DCs, along with the control, remained below 100Å^2^ throughout the MDS trajectories (**Fig. 6B**). The Polar Surface Area (PSA) of a compound represents the surface area attributed to polar atoms, such as oxygen and nitrogen, along with their attached hydrogen atoms. The PSA values for DC1 – DC4 were recorded in the 150Å^2^ range, although DC3 initially showed a lower value. Meanwhile, DC6 and DC4 exhibited PSA values of 120Å^2^ and 100Å^2^, respectively. Compared to the control compound, fluctuations were observed in the PSA curve, though the values remained close to those of the DC1 – DC4 compounds (**Fig. 6C**). The Radius of Gyration (rGyr) measures the compactness or distribution of a DC’s atoms around its center of mass. By tracking rGyr over time during the simulation, conformational changes in the DCs were observed. The highest conformational displacement was recorded for DC2, DC1, and DC3. In contrast, the conformational displacement and mass distribution for the other compounds, as well as the control, displayed stable rGyr plots throughout the simulation period (**Fig. 6D**)

**Fig. 6.**
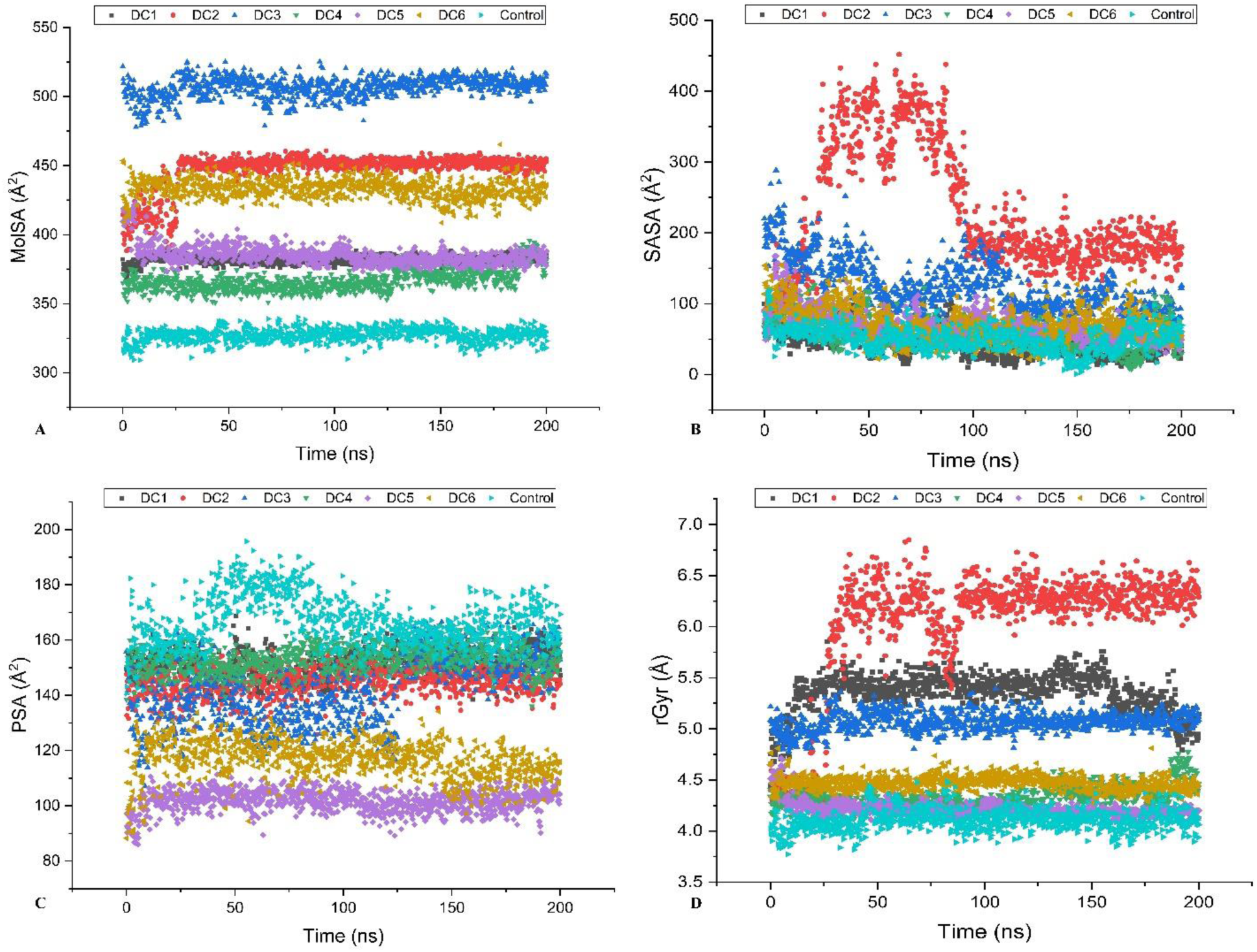
Graphical representation of four key chemical properties of DCs, focusing on surface area, solvent exposure, polarity, and extendedness. The parameters (A) Molecular Surface Area (MolSA), (B) Solvent Accessible Surface Area (SASA), (C) Polar Surface Area (PSA), and (D) Radius of Gyration (rGyr) were calculated over the 200 ns simulation and are depicted in different colors on the plots, illustrating the variations in these properties throughout the simulation.

### Free energy calculation reveals potential strengths of interaction between DCs and DfrA1

One of the key aspects in drug discovery is the calculation of the MM/GBSA status, which is widely used to assess the stability and affinity of ligand binding during the drug discovery process. The calculated free energy for the r_psp_MMGBSA_dG_Bind_vdW module was ≤ −60 for all DCs except the control, indicating that DCs are more fit to the receptor than the control drug, Iclaprim. Positive values were reported for r_psp_MMGBSA_dG_Bind_Solv_GB for both the DCs and the control, suggesting that the binding of the DCs would not readily be influenced by the solvation environment. The reported negative values of ≤ −40 for r_psp_MMGBSA_dG_Bind_Lipo indicate the contribution of non-polar solvation energy (lipophilic interactions) to the binding free energy in MM/GBSA calculations. Subsequently, r_psp_MMGBSA_dG_Bind_Coulomb measured the contribution of Coulombic (electrostatic) interactions to the binding free energy in MM/GBSA calculations (**Fig. 7**). The consistently negative free energy values for this parameter across all DCs highlight the role of Coulombic forces in the binding affinity between the DCs and DfrA1. The final module of MM/GBSA, r_psp_MMGBSA_dG_Bind, was used to estimate the binding strength of DCs to the DfrA1 binding pocket. The ΔG Bind values for all compounds were higher than those for the control, with values ≤ −90, indicating a stronger and more stable binding to the targeted residues compared to the available drug (e.g., Iclaprim) (**Fig. 7**).

**Fig. 7.**
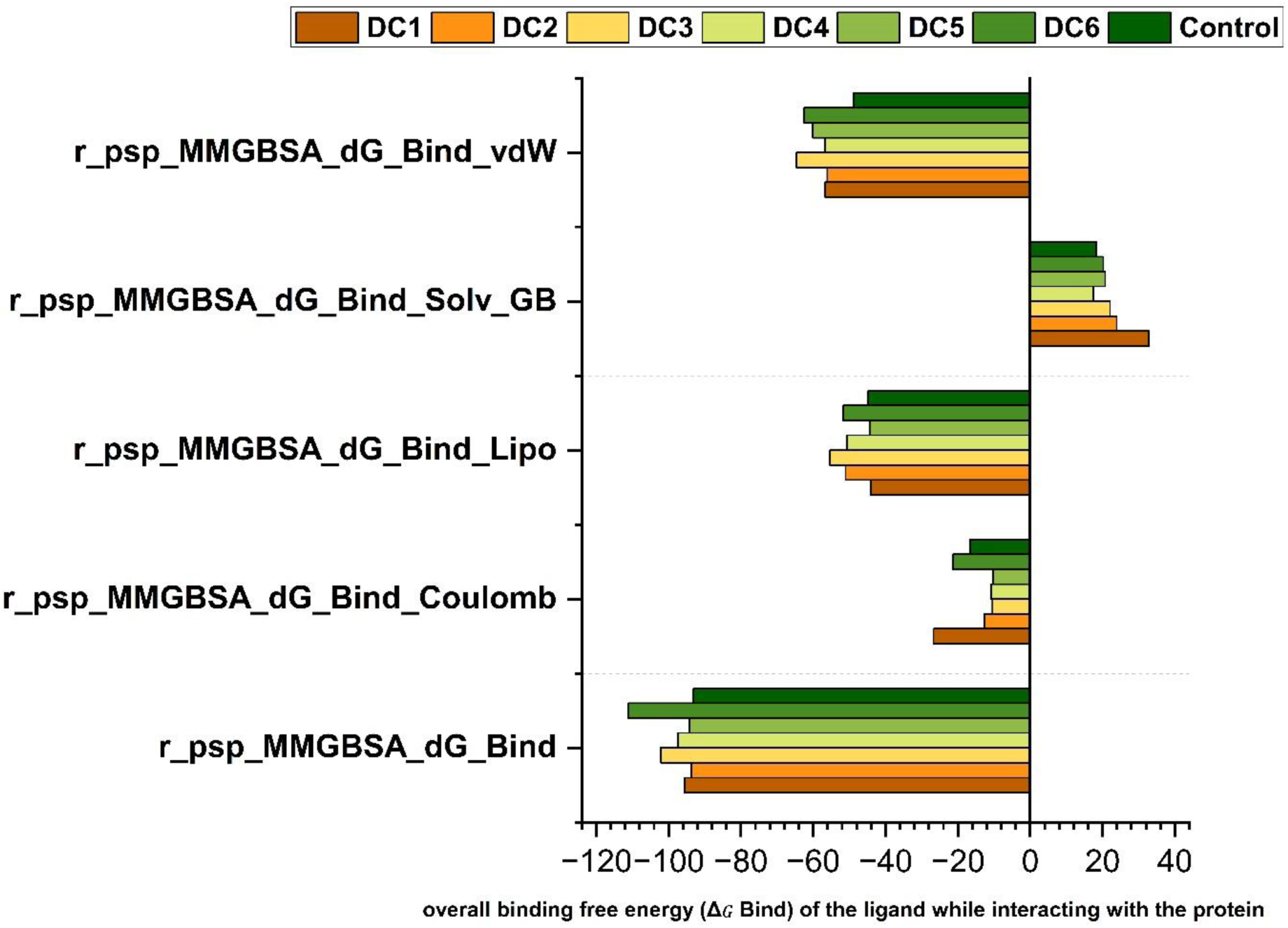
Validation of thermal MM-GBSA analysis for six drug candidates (DCs) and control while interacting with DfrA1. Total MM-GBSA binding free energy was calculated, considering binding free energy, generalized Born electrostatic solvation energy, lipophilic effects, electrostatic energy (Coulombic effect), and total release free energy during interaction with the pocket residues of DfrA1. A bar graph with distinct colors represents each compound included in the study.

## Discussion

Trimethoprim (TMP) resistance in bacterial pathogens significantly limits treatment options, leading to more severe infections, particularly with *E. coli*, *P. aeruginosa* and *K. pneumoniae* ^7,9,11^. The spread of resistance genes exacerbates this issue, increasing healthcare costs, prolonging illness, and posing a growing public health threat due to rising multidrug-resistant organisms. Thus, computational drug discovery approach may overcome TMP-resistance by employing structure-based design, virtual screening, *de-novo* drug design, and MDS profiling to identify novel, effective, and safer antibacterial therapeutics ^24,25^. In this study, we evaluated 3,601 newly synthesized chemical compounds against the TMP-resistant DfrA1 protein using supercomputer-based optimization and screening to identify novel drug candidate(s). This research marks a significant step toward addressing AMR, particularly TMP resistance in bacterial pathogens like *E. coli*, *P. aeruginosa*, and *K. pneumoniae*, offering promising therapeutic solutions for combating infections caused by these superbugs. This innovative approach utilized advanced drug discovery tools for precise compound interaction analysis and semiempirical quantum mechanical methods to refine leading candidates’ structures, enhancing their biological activity and paving the way for more targeted and effective therapeutic solutions ^26^. The use of a cluster-based supercomputer greatly expedited the optimization process.

A major breakthrough of this study was the identification of six novel inhibitors (DC1 – DC6) targeting the TMP-resistant DfrA1 protein from a library of 3,601 optimized compounds. These top candidates were selected for their robust H-bonding and non-covalent interactions, which are critical criteria in the drug discovery process ^27,28^. These six DCs with strong H-bonding and lower binding free energies demonstrated favorable interactions with the target protein, DfrA1 compared to the control drug, Iclaprim. Moreover, DFT calculations for the six DCs revealed that DC6 had the lowest HOMO-LUMO energy gap (3.18 eV), indicating that DC6 is more reactive than the other DCs. The HOMO-LUMO energy gap of Iclaprim was measured at 4.77 eV, closely aligning with the ranges of 4.41 to 5.46 eV observed for the other five DCs. Moreover, UV-Vis calculations evaluated the absorbance characteristics of the DCs, showing that DC6 exhibited the highest epsilon value (absorption coefficient) in comparison to the other candidates and the control drug. Besides, DC1, DC2, and DC3 exhibited absorption peaks at shorter wavelengths than the control, indicating lower excitation energy and enhanced reactivity, thereby suggesting their potential effectiveness as therapeutic agents. The lower excitation energy at these wavelengths suggests increased reactivity for these compounds, indicating their potential effectiveness as therapeutic agents ^29,30^. The next phase of investigation focused on the ESP of the DCs. The higher ESP values observed for the DCs, relative to the control, suggest that these compounds possess highly interactive electrostatic surfaces, enhancing their potential for molecular interactions ^31^. This characteristic enhances their potential as effective inhibitors of DfrA1, indicating that their electrostatic properties may promote stronger interactions with the target protein.

While the initial screening suggested strong interactions among six DCs with the DfrA1 protein, MDS revealed that only DC1, DC4, and DC6 maintained stable interactions, whereas DC2 exhibited no significant hydrogen bonding with the DfrA1 pocket residues. This stability and higher interaction fractions for DC1, DC4, and DC6 suggest their greater potential as effective inhibitors of DfrA1 (**Fig. 4**). Interestingly, the RMSD plots of DfrA1 while interacting with DCs showed fluctuations within a range of 1-3 Å, which is perfectly acceptable ^32^. Notably, DfrA1 exhibited fewer displacements in the RMSD plots during interactions with the DCs compared to the control drug. Among these DCs, DC2 displayed the highest fluctuations in the RMSD plot. Additionally, the UV-Vis analysis revealed that DC2 exhibited unusual absorption characteristics, likely contributing to its instability. In contrast, only DC4 and DC6 demonstrated excellent RMSD values, remaining below 2 Å, which indicates strong and stable interactions with the DfrA1 binding pocket. This finding aligns with previous studies in computational drug chemistry, suggesting that such low RMSD values are indicative of reliable binding stability ^33,34^. The analysis of four key parameters namely MolSA, SASA, PSA, and rGYr revealed that DC4 and DC6 were the most effective inhibitors of DfrA1, with values comparable to or better than the control drug. Furthermore, the MM/GBSA calculations supported these findings by evaluating the overall binding free energy (ΔG Bind) of DC4 and DC6 during their interactions with DfrA1. One of the critical issues for TMP resistance against DfrA1 protein is the amino acid substitutions at D27E and L28Q position ^23^. While Iclaprim is effective against mutated DfrA1 because of its capacity to interact with the altered amino acids, the organofluorinated compound DC4 and the benzimidazole compound DC6 showed notable advantages over Iclaprim in terms of thermochemical behavior, molecular dynamics, and binding affinities. Benzimidazole and organofluorinated compounds are potent antibacterial agents commonly employed to combat infections caused by Gram-negative and certain Gram-positive bacteria ^35,36^. This suggests that DC4 and DC6 could be effective treatments against the TMP-resistant DfrA1 protein. Their promising performance highlights their potential as viable alternatives in the fight against resistant bacterial infections. Since the findings of this study are based solely on *in-silico* computational methods, further wet lab experiments for clinical verification and molecular functional characterization are essential to validate the findings. These steps are essential for confirming new drug candidates against TMP-resistant DfrA1 before considering their clinical applications.

## Conclusion

Antibiotics have long been the cornerstone of combating bacterial infections, yet the growing challenge of AMR underscores the critical need for alternative therapeutic options. This study aimed to identify potential inhibitors targeting the TMP-resistant DfrA1 protein, which contributes to development of AMR in a wide range of bacterial pathogens including *E. coli*, *K. pneumoniae* and *P. aeruginosa* both in humans and animals. Our high-throughput computational analysis reveals the efficacy of DC4 and DC6 as potential inhibitors of DfrA1, highlighting their strong inhibitory affinities. These findings suggest DC4 and DC6 as promising candidates in the battle against DfrA1-carrying resistant pathogens. While the findings of this study pave the way for discovering novel therapeutic candidates against TMP-resistant bacterial pathogens, further validation through real-world clinical experiments is essential.

## Method and materials

### Compounds refinement

A total of 3,601 newly synthesized chemical compounds were extracted from [ChemDiv](https://www.chemdiv.com/) database ^37^. We ensured that the compounds maintained the Lipinski’s rules of 5 during extraction. The initial structures were converted into molecular data files (.MOL) using the structure transformation tool in MarvinSketch v24.1.3 ^38^. Subsequently, all compounds underwent preliminary geometry optimization through the Optimize Geometry module in Avogadro ^39^. To enhance compatibility with downstream analysis software, the files were converted to the PDB using Avogadro’s program extension tool. For further refinement, these molecules were subjected to optimization using Gaussian 09W (Gaussian, Inc., Wallingford CT, 2016) with the Universal Force Field (UFF) method, simulating a water solvent environment ^40^. This optimization was performed on a supermicro-built, server-based supercomputer, significantly reducing processing time and facilitating efficient handling of the large dataset.

### Protein system configuration

The crystallographic structure of the DfrA1 protein (PDB ID: 5ECX), extracted from *K. pneumoniae* and expressed in *E. coli*. DfrA1 is linked to TMP resistance and has been experimentally validated, was retrieved from the RCSB Protein Data Bank^41^. The three-dimensional (3D) structure was resolved using X-ray diffraction at a resolution of 1.95 Å. The DfrA1 structure was meticulously refined using Maestro v2024-4 (Schrödinger, LLC, New York, NY, 2024), which ensured the removal of undesirable interactions and optimized the engagement between potential binding compounds and the target protein during the molecular screening process ^42^. For the refinement, the protein preparation was conducted with the protein preparation workflow of the Maestro with OPLS_2005 force field ^43^. During the refinement process, missing side chains were added, hydrogen (H)-bonds were optimized, and excess water molecules were removed from the protein structures.

### Binding simulation of protein and compounds

Prior to performing binding analysis, the SiteMap module of Maestro (Schrödinger, LLC) was utilized to determine the drug-binding side regions in the targeted DfrA1 protein ^42^. The binding simulation between the DfrA1 and compounds was performed using the vina.bat scripts executed via PowerShell for Linux ^44^. The configured protein structure was recognized as a biological target by the system, with parameters such as the center and dimensions included to create an optimal environment. The system’s input energy was set to 64, and the exhaustiveness was configured to 128 to reduce the likelihood of false interactions between the protein and binding compounds. This approach enabled the simultaneous analysis of the binding efficiency of 3,601 compounds, streamlining the process and reducing complexity. Finally, six DCs such as DC1 – DC6) that demonstrated strong chemical interactions, including H-bonding and non-covalent forces, with DfrA1 were selected for advanced analysis based on their proven effectiveness and promising results.

### Orbital energy analysis

The calculations of HOMO and LUMO were conducted using Gaussian 09W (Gaussian, Inc.) ^45^. Assessing these frontier molecular orbitals (FMOs) was crucial for understanding the chemical reactivity of the DCs. DFT was employed to determine the electronic structure, utilizing the B3LYP method and the 6-31g (d,p) basis set, with Opt+Freq analysis modules chosen for the calculation process under gas phase conditions to evaluate HOMO-LUMO gaps ^46^. All the processors of the cluster-based supercomputer were utilized to perform the calculations. The energy gap calculation plots were generated using GaussSum ^47^.

### Electrostatic surface potential and UV-Vis spectra of drug candidates

The electrostatic surface potential (ESP) and UV-Vis spectra for each of the six DCs, along with the control drug Iclaprim, were determined using Gaussian 09W (Gaussian, Inc.). For the ESP analysis, the surface-contours module of Gaussian was utilized with the addition of a new cube action for the selected molecules. Initially, a cube was generated to calculate the total electron density of each compound using the SCF density matrix and a coarse grid with a point value of - 2. After calculating the electron density, the ESP surface was analyzed based on the resulting molecular surfaces ^48^. UV-Vis spectra were obtained using the vibrational spectra module in Gaussian (Gaussian, Inc.). The IR spectrum data, in a text-readable format, were exported from the UV-Vis plot and further analyzed using GraphPad Prism version 10 for Windows (GraphPad Software, San Diego, CA, USA).

### Molecular dynamics modeling

To simulate the interactions between the DCs and DfrA1 (the biological target), we selected complexes that exhibited effective binding during the binding simulation analysis. These complexes were then prepared and subjected to energy minimization using the Protein Preparation Wizard, where missing side chains and loops were added using Prime (Schrödinger, LLC). Molecular dynamics simulations were then carried out using the Desmond MD engine ^49^. The systems were solvated in a cubic box with a 10 Å edge length using the SPC water model, and ions were added to ensure a neutral net charge. The final setups included approximately 21,000 atoms. Prior to starting the production simulations, the systems underwent the standard relaxation protocol provided by Desmond. Production simulations were performed under an NPT ensemble, with pressure maintained at 1.01325 bar using the Martyna–Tobias–Klein method and temperature regulated at 300 K with the Nosé–Hoover thermostat ^50^. A Coulombic cutoff radius of 9 Å was used, following the default settings. Each system was simulated for a duration of 200 ns.

### Quantification and analysis of the drug candidate surface properties during simulation

The solvent accessible surface area (SASA) (Å^2^), polar surface area (PSA) (Å^2^), molecular surface area (MolSA) (Å^2^), and radius of gyration (rGyr) (Å) of the DCs were meticulously quantified utilizing custom simulation.py scripts provided by Maestro (Schrödinger LLC) ^42^. These calculations were conducted at 1 ns interval throughout the simulation, ensuring detailed temporal resolution. The surface dynamics of the compounds were then extracted from the trajectory files, resulting in a comprehensive dataset for further analysis. To facilitate clear and insightful visualization, the results were processed and graphically illustrated using OriginPro 2024 (OriginPro software, Northampton, USA) on a Windows platform, allowing for an in-depth exploration of the DCs surface properties over the course of the simulation.

### Biding free energy calculation of the drug candidates

The binding free energy for the DCs interacting with the target protein DfrA1 was calculated using the MM/GBSA method, implemented via Maestro (Schrödinger LLC) ^42^. Following a comprehensive 200 ns simulation, individual DfrA1 and DC structures were extracted from the simulation trajectory. These extracted structures were then input into the MM/GBSA framework to compute the free energy of interaction ^50^. The calculation employed the novel variable dielectric surface generalized born (VSGB) solvation model, optimized for high-resolution protein structures, to ensure accurate estimation of solvation effects ^51^.

## Data availability

All data generated or analysed during this study are included in this published article [and its supplementary information files]

## Author contributions

S.H., M.M.M., M.N.H., and T.I. conceived and designed the study; S.H., S.R., M.B.A., F.M.S., T.S., N.T.S., S.S., F.Y., A.C., and A.D.A.S. were responsible for data acquisition, analysis, and writing the original draft. M.M.M., M.N.H., and T.I. conducted the critical review and editing. All authors contributed to the article and approved the final submitted version.

## Supplementary Information

**Fig. S1**. Binding pocket of DrfA1.

**Fig. S2**. UV-Visible spectra of selected drug candidates.

**File S1**. Docking scores and RMSD values of 3,601 chemical compounds.

**Table S1**. Selected vibrational frequencies (cm^-1^) of drug candidates (DCs) calculated in gas phase (scaled).

